# Origin, Trophic Transfer And Recycling Of Particulate Organic Matter In The Waters Of Two Upwelling Bays Of Humboldt Current System: Insights From Compound-Specific Isotopic Compositions Of Amino Acids

**DOI:** 10.1101/2024.06.24.600486

**Authors:** B.M Srain, J Valdés, A Camaño

## Abstract

Particulate organic matter (POM) is considered the primary source of N and C in the ocean. In pelagic marine environments, POM consists of algae and detrital nitrogen, with amino acids representing the largest chemical fraction. Currently, measurements of the isotopic distributions of N atoms in amino acids are considered powerful tools for exploring and determining the metabolic sources involved in the synthesis and degradation of organic matter. In this study, we measured the δ^15^N of amino acid signatures (δ^15^N-AA) in suspended and sinking POM collected from two upwelling bays in northern Chile, to examine isotopic enrichment patterns and gain insights into the origins, trophic transfer, and heterotrophic reworking of this organic fraction. At Mejillones Bay, the δ^15^N-AA values of suspended POM ranged from 5 ‰ to 27 ‰, while at Antofagasta Bay, these values oscillated between 9 ‰ and 24 ‰. The sinking POM collected from sediment traps exhibited values and isotopic fractionation patterns similar to those observed in the deeper layers of the water columns in both bays. The enrichment patterns of δ^15^N-phenylalanine and δ^15^N-NO_3_ ^-^ demonstrated the autochthonous character of the POM and its predominantly marine origin at both bays. The parameters trophic transfer (ΔTr) and heterotrophic reworking (ΣV) indicated that the heterotrophic recycling of POM occurs more intensively at through the oxyclines. Furthermore, these parameters revealed an enhanced trophic transfer magnitude and higher heterotrophic re-synthesis of POM in the waters of Mejillones Bay, resulting in a lower flux of exported POM than that observed in Antofagasta Bay. These differences highlight the spatial heterogeneous nature of organic matter transfer and reworking processes in this upwelling system.

## 1. INTRODUCTION

Most organic matter in natural environments exists not in living organisms but as detrital nitrogenous material in particulate organic matter (POM) (1). Consequently, POM is considered the main source of nitrogen, carbon, and other biologically important ocean elements, supporting pelagic and benthic ecosystems (2). The production, alteration, and degradation of POM are fundamental processes in the biogeochemical cycles of carbon and nitrogen across various environments, including oceanic and coastal waters (3,4), marine sediments (5), and soils (6). In pelagic marine environments, suspended and sinking particles contain algae and detrital nitrogen, with amino acids (AA) representing the largest fraction of organic nitrogen exported to the ocean’s interior (7,8). The isotopic fractionation of carbon and nitrogen in AA provides a direct record of central metabolic cycles (9), sources, trophic transfers, and heterotrophic microbial reworking of organic matter (10–13). Therefore, measurements of the isotopic distributions of carbon (14,15) and nitrogen (16) atoms in AA are currently considered powerful tools for exploring and determining the metabolic sources involved in the synthesis of organic matter, as well as the transformation processes during its decomposition through autotrophic and heterotrophic metabolism (1,17–19). Furthermore, δ^15^N patterns in AA offer exceptionally detailed tracers for understanding trophic changes (16). Given that each AA is synthesized through distinct biosynthetic pathways in both autotrophs and heterotrophs, δ^15^N-AA patterns can provide robust indicators of nitrogen sources and mechanisms of organic matter alteration (10). Under this conceptual framework, δ^15^N-AA data indicators have been developed to track a wide range of processes in detrital organic matter, including estimates of trophic transfers (20,21), the relative degree of intracellular heterotrophic bacterial resynthesis (e.g., parameter “ΣV”) (1), extracellular heterotrophic bacterial degradation (22,23), as well as estimates direct measurements of the δ^15^N-AA value, to determine the inorganic sources of N for a system (24–26).

The Mejillones Peninsula (*ca*. 23°S – 70°W; Fig. 1), is the most prominent geographical feature of tectonic origin on the northern coast of Chile. It extends 50 km in length and 20 km in width, comprising three morphological units: two north-south mountain ranges separated by a coastal plain (27). This peninsula disrupts the coastline’s linearity, creating two bays with opposite orientations: Mejillones Bay, oriented towards the equator (north), and Antofagasta Bay, facing south (28–30). This area is part of the Humboldt Current System, one of the world’s most productive ecosystems (31,32). The oceanographic conditions of Mejillones Bay are influenced by forcings that operate at various spatiotemporal scales. Coastal upwelling, driven by wind action, brings Subsurface Equatorial Water, rich in nutrients and low in oxygen, to the surface. This water frequently enters the bay, creating natural hypoxic or suboxic conditions (28,33,34). The bay’s northern orientation causes upwelling shadows, primarily determined by the coastline’s morphology. This leads to the formation of a thermal front at the bay’s mouth, promoting stratification and retaining water within the bay (27,28,30,33). The alternation of vertical mixing and stratification processes in the water column and the presence of upwelling shadows are crucial for organism retention and the generation of high primary productivity and organic matter within the bay (32,35).

**Fig 1.**
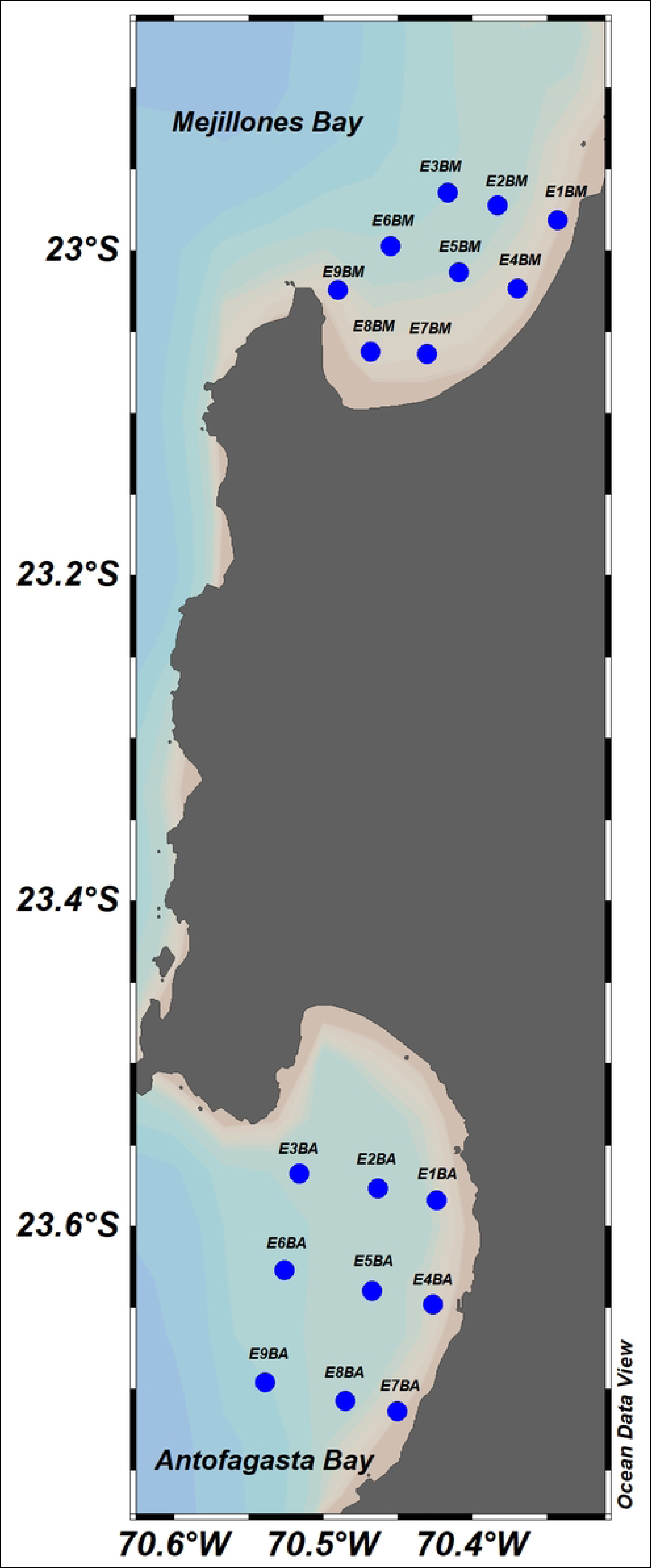
Geographical location of the study site at the bay system of the Mejillones Peninsula in northern Chile.

In contrast, Antofagasta Bay is directly influenced by the Humboldt Current system (29,36). The hydrodynamics within the bay are shaped by a thermal front created by an upwelling center in the southern part, which regulates the bay’s filling and emptying processes. This model highlights the entry of water through the peninsular sector and its exit through the southeastern sector, generating a cyclonic turn in the bay’s center (37,38). The development of thermal fronts in Antofagasta Bay has been described as physical barriers typical of upwelling zones, causing a retention of water masses towards the coast, a phenomenon known as the “upwelling trap” (29,30). This trap is efficient in retaining organic and inorganic particles and planktonic organisms (38). The hydrodynamic conditions of Antofagasta Bay are characterized by low-intensity surface currents compared to the oceanic zone. During summer, the main flow brings water into Antofagasta Bay, while during winter, in the absence of winds, the opposite occurs (29,30,37,39,40). Unlike Mejillones Bay, Antofagasta Bay has seen limited research efforts to understand its functioning. A few studies have mainly focused on oceanographic aspects such as temperature variability patterns, seasonal dynamics of zooplankton, phytoplankton ecology, the coupling of physical and biological processes, and the hydrodynamics within the bay. In this study, we analyzed the δ^15^N-AA signatures in suspended and sinking POM collected from two upwelling bays in northern Chile. Our aim was to examine the enrichment patterns and behaviors of δ^15^N-AA parameters to determine the origins, sources, and mechanisms of trophic transfer and heterotrophic microbial reworking of POM through the water columns.

## 2. METHODS

### 2.1. Sampling

Nine subtidal stations were sampled at Mejillones and Antofagasta Bays in March 2023 (Fig 1). Water samples were collected from four depths (5, 20, 35, and 45 m) using Niskin bottles (10 L) and stored in acid-washed carboys in the dark at approximately 10 °C. Once in the laboratory, 30 L of seawater from each depth were filtered through pre-combusted 0.7 μm glass fiber filters (Whatman GF/F). The filtrates and filters were kept at −80 °C before analyzing total organic carbon (TOC), chlorophyll-a, chlorines, and nutrients (NH_4_ ^+^, NO_2_ ^-^, NO_3_ ^-^, and PO ^-^). Continuous profiles of temperature (°C), salinity, dissolved oxygen (mg/L), and pH were measured using a CTD-O (SBE-19) instrument.

In each bay, three special stations were established, oriented north-south along the central axes of the bays (Stations E2BM, E5BM, E7BM, E2BA, E5BA, and E8BA; Fig. 1). Water samples were collected from four discrete depths (5, 20, 35, and 45 m) for the analysis of δ^15^N compound specific isotope analyses of AA on POM and specific water samples for δ^15^NO_3_ ^-^ analysis. These subsamples were collected into 20 mL acid-washed plastic scintillation vials using a syringe with a GF/F filter. Additionally, sediment traps were placed 5 m above the ocean floor.

### 2.2. Nutrients

Water samples were stored in clean plastic bottles and kept in the dark at −20 °C until analysis by spectrophotometry in the laboratory for dissolved inorganic nutrients analysis. The dissolved inorganic nutrients NH_4_ ^+^, NO_2_ ^-^, NO_3_ ^-^, PO_4_ ^3-^, and Si(OH) were analyzed in the laboratory using UV-VIS spectrometry (41,42).

### 2.3. Chlorines

The suspended particulate material captured in a 0.7 μm glass fiber filter was mixed with 5 mL of 90% acetone in 15 mL polypropylene centrifuge tubes for chlorine analyses.

Samples were homogenized using a vortex mixer and sonicated for 10 min. The liquid phase (supernatant) was recovered after centrifugation at 3000 rpm for 5 min and transferred to 20 mL vials (3x). Samples were analyzed using a Turner Design 10.005 fluorometer after dilution, before, and after acidification with 10% HCl, using the appropriate pigment standard (Sigma-Aldrich C6144-1MG) following the standard technique described by Holm-Hansen et al. (43). The chlorine concentration was calculated as the sum of the concentrations of chlorophyll-*a* and phaeopigments.

### 2.4. Total organic carbon

For TOC analysis, the water samples were homogenized and diluted, and a small aliquot was injected into a heated reaction chamber packed with a platinum catalyst to vaporize the water and oxidize the organic carbon compounds to CO_2_ and H_2_O. The CO_2_ from the oxidation of organic carbon was measured by nondispersive infrared detection using an elementary analyzer.

### 2.5. Isotopic analysis

#### 2.5.1. Amino acid hydrolysis and isotopic analysis

All samples were hydrolyzed and derivatized as previously described (1,44). Dried samples were hydrolyzed under standard conditions (6 N HCl for 20 h at 110 °C), and the resulting hydrolysate purified using cation exchange chromatography (Dowex 50WX8-400 ion exchange resin) (42). Isopropyl-TFA derivatives were prepared as described by Silfer et al. (46). Derivatized samples were analyzed by a Thermo Trace 1310 gas chromatograph with IsolinkII/ConfloIV (reactor 1000 °C), coupled to a Thermo Delta V isotope ratio mass spectrometer.

AA were separated for δ^15^N analyses using a BPX5 column (60 m × 0.32 mm, 1 μm film thickness; SGE Analytical Science, Trajan, Austin, TX, USA) and for δ^13^C analyses using a DB-5 column (50 m × 0.32 mm 0.52 μm film thickness; Agilent Technologies, Santa Clara, CA, USA). Under our analytical conditions, both δ^15^N and δ^13^C values could be reproducibly measured for alanine (Ala), aspartic acid + asparagine (Asp), glutamic acid + glutamine (Glu), leucine (Leu), isoleucine (Ile), proline (Pro), valine (Val), glycine (Gly), lysine (Lys), serine (Ser), phenylalanine (Phe), threonine (Thr), and tyrosine (Tyr). The reproducibility of these AA was typically less than 1‰. The amino acid δ^13^C values were determined from the measured values of the AA derivatives following the approach of Silfer et al. (46), with corrections based on an AA mixture standard for which isotopic values had been independently determined by offline elemental analyzer analysis. The directly measured δ^15^N-AA values were also corrected based on bracketing external standards, as described in McCarthy et al. (44).

The injector temperature was 250 °C with a split He flow rate of 2 mL/min. The GC temperature program for nitrogen isotope analysis was: initial temp = 70 °C hold for 1 min; ramp 1=10 °C/min to 185 °C, hold for 2 min; ramp 2 = 2 °C/min to 200 °C, hold for 10 min; ramp 3 = 30 °C/min to 300 °C, hold for 6 min. The GC temperature program for carbon isotope analysis was initial temp = 75 °C hold for 2 min; ramp 1 = 4 °C/min to 90 °C, hold for 4 min; ramp 2 = 4 °C/min to 185 °C, hold for 5 min; ramp 3 = 10 °C/min to 250 °C, hold for 2 min; ramp 4 = 20 °C/min to 300 °C, hold for 5 min.

#### 2.5.2. δ^15^N of dissolved nitrate (δ^15^NO_3_ ^-^)

The stable isotopic composition of the naturally abundant N in NO_3_ ^-^ was analyzed using denitrification (47,48). The samples were previously treated with sulfamic acid to reduce and eliminate the naturally present NO_2_ ^-^ (49,50). δ ^15^N values were measured in N_2_O quantitatively produced by denitrifying bacteria that lack N_2_O-reductase activity. The N_2_O produced was analyzed using GC CF-IRMS (Finnigan Delta Plus). Data are presented in delta notation as δ^15^N = (R (sample)/ R (reference) – 1) x 1000/, where R = ^15^N/^14^N. The isotopic reproducibility was greater than 0.2%.

### 2.6. δ^15^N AA parameters for N source, trophic transfer, and resynthesis

The ΔTr value, which indicates the specific number of trophic transfers, was calculated following McCarthy et al. (1), defining the average offset between these same AA in phytoplankton, as ΔTr = 0, and then using the corresponding δ^15^N for a single trophic transfer, as reported in the culture experiments (food-consumer = 4.67) (16).

Thus:

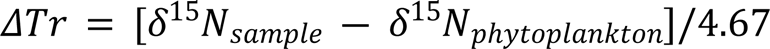

Where:

δ^15^N sample is the average δ^15^N of trophic AA Asp, Glu, Leu, Val, and Pro. The δ^15^N phytoplankton is the average value of the source AA Gly, Phe and Lys.

The parameter Σ*V*, which indicates total heterotrophic reworking, is defined as the average deviation in the δ^15^N values of the trophic amino acids Ala, Asp, Glu, Ile, Leu and Pro (51).

Thus:

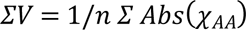

Where:

“χ” is the deviation of each of the trophic AA with respect to the average of all trophic AA. “n” is the total number of trophic AA used in the calculation

In this study, we use the term “heterotrophic resynthesis” as the heterotrophic reworking of proteinaceous material, mediated by multiple processes carried out by planktonic heterotrophic organisms, such as bacteria, archaea, protists and zooplankton, including hydrolysis, uptake and de novo synthesis, salvage AA incorporation into new protein, as well as strict catabolism, as was defined by McCarthy et al. (1)

The δ^15^-N and δ^13^C of total hydrolyzable AA (THAA), was estimated by averaging the δ^15^- N values of each specific AA and whose values were operationally considered here as the bulk δ^15^-N values for suspended and sinking POM, assuming the bulk δ^15^-N estimations, as the integration of all nitrogen-containing entities in our POM samples (10).

### 2.7. Amino acid grouping

According to the behavior of individual δ^15^N values within food webs (16), AA can be classified into two groups:

i. The trophic AA Asp, Glu, Ala, Ile, Leu, Val, and Pro were strongly enriched with trophic transfer.
ii. Source AA: Gly, Thr, Phe, Tyr, and Lys, whose δ^15^N values remain largely unaltered with each trophic shift.

Since Phe has been reported and described as an abundant AA and is considered the most stable reference source AA in published culture experiments (1,16), we will focus our discussion on the origin and sources of suspended and sinking POM in waters of the bays of Mejillones and Antofagasta, only based on the δ^15^N-Phe values.

### 2.8. Ancillary oceanographic parameters

Dissolved inorganic N anomalies, defined as a linear combination of NO_3_ ^-^ and PO_4_ ^3-^, were calculated to determine the roles and distribution of N fixation and denitrification in the water column. The parameter N^∗^ was estimated following Hansell et al. (52) (N∗ = [NO_3_^−^ + NO_2_ ^−^ + NH ^+^] – 16 [PO_4_ ^3-^] + 2.9). N^∗^ values lower than −3 μmol kg^−1^ are indicative of denitrification while values higher than 2 μmol kg^−1^ denote nitrogen fixation (53).

The apparent oxygen utilization (AUO), which indicates the amount of oxygen respired in the ocean interior, was calculated as follows:

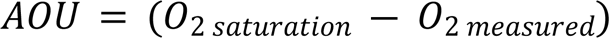

Oxygen saturation as a function of temperature and salinity in the water column was determined according to Weiss, (54).

### 2.9 Dissolved oxygen inventories and water column stability

Oxygen inventories in the water column were calculated integrating the vertical profiles of dissolved oxygen using the Sigma Plot 10.0 software. The stability of the water column was estimated by calculating the Brunt Väisälä Frequency using the Ocean Data View software, version 5.6.4 (138).

### 2.10. Sediment traps deployment

During the cruise, five surface-tethered sediment traps were deployed and attached to a 70 kg weight at stations E5BM, E7BM, E2BA, E5BA, and E8BA (Fig. 1). The traps were submerged at a depth of 45 m for 30 days to collect suspended particulate organic matter (POM) that passively sinks through the water columns of the bays, allowing for the determination of the downward flux of particles. The setup consisted of a stainless-steel cross supporting four individual PVC pipe collectors, each equipped at the base with stopcocks coupled to particle interceptor traps (PITs) made of four 1 L high-density polyethylene bottles. As a control, one bottle was kept with the stopcock closed during deployment. Following the JGOFS protocol (55), each PIT was filled with 900 mL of filtered seawater brine solution (50 g of NaCl per liter of seawater) and 100 µL of 0.1% HgCl_2_ solution to inhibit biological activity. The resulting density gradient allowed particles collected through the tubes to enter the PITs while preventing their escape due to reflux and washing during trap removal. The trap arrays had an aspect ratio of 10.5. Upon trap retrieval, the PITs were immediately removed and stored in the dark at 4 °C until laboratory analysis. The samples contained in the PITs were filtered using pre-combusted fiberglass filters with a 47 mm diameter and 0.7 µm pore size to retain the particulate matter. The filters were subsequently freeze-dried, weighed, and stored at −20 °C until chemical analysis. The same procedure was used for the control samples.

### 2.11. Data analysis

Homogeneity of variances (Levene test) and normality of variables (Shapiro–Wilk test) were not fulfilled; therefore, we tested for significant differences using the non-parametric Mann-Whitney and Wilcoxon tests. Correlations were examined using Spearman R coefficients. All statistical analyses were performed using the package software Statistica version 12.0.

## 3. RESULTS

### 3.1. Hydrography

The vertical physical-chemical structures of the water columns of Mejillones Bay and San Jorge Bay were virtually similar, showing the expected patterns for the area and season during which this study was carried out (Figs 2 A-J). In both bays, a warmer, less saline and completely oxygenated surface layer (0-5 m) was evident with temperatures ranging between 14 °C and 22 °C, salinities between 33.5 and 34.5 PSU and dissolved oxygen between 4 and 10 mg/L (Figs 2 A-C, F, G and H). From 5 m to 40 m depth, the development of a marked thermocline and oxycline was evident, where the temperature constantly decreased from the surface to the bottom, dropping up to 13.2 °C and a minimum dissolved oxygen content of 0.4 mg/L, denoting the occurrence of hypoxic and suboxic waters inside both bays. The halocline was noticeably shallower, lying between the surface and a depth of 5 m, in both bays (Figs 2 B and G). Starting at a depth of 5 m and towards the bottom, the salinity showed a quasi-homogeneous vertical structure with an average value of 35.1 PSU (Figs 2 B and G). Both in Mejillones and San Jorge bays, more alkaline waters were observed in the first 12 m, with pH values that oscillated between 8 and 8.6 pH units (Figs 2 D and I). Below 20 m, the water became more acidic, with pH values that decreased to 7.8 pH units (Figs 2 D and I). The vertical physicochemical structure of the water columns in both bays suggested at least two masses of water inside them or a mixture of them. As indicated by the T-S diagrams (Figs 2 E and J), both bays contained Subtropical Surface Water (SSTW) and modified Equatorial Subsurface Water (AESS; Figs 2 E and J).

**Fig 2.**
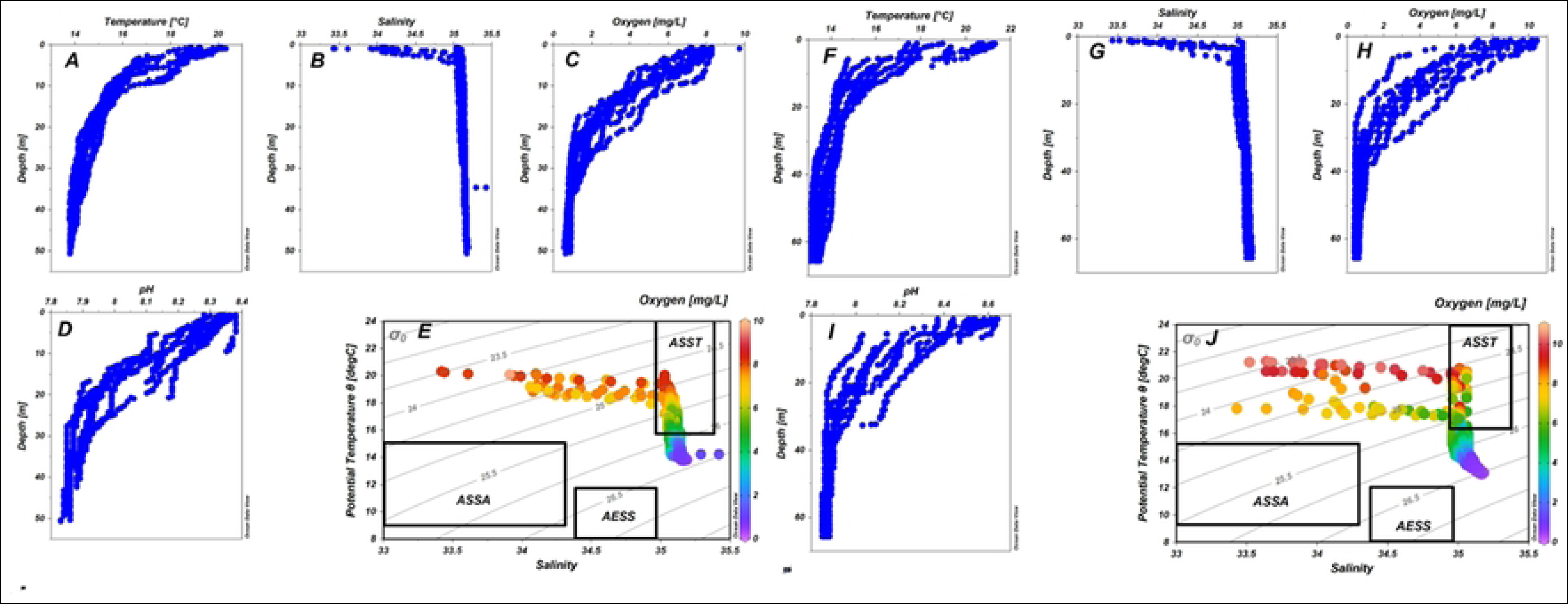
Vertical profiles of temperature (°C), salinity, dissolved oxygen (mg/L), pH, and T-S diagrams for Mejillones Bay (A-E), and Antofagasta Bay (F-J).

### 3.2. Chlorophyll-*a*, and nutrient contents

Canonical vertical profile distribution of chlorophyll-*a* was similar in both Bays, with the highest content occurring in the most superficial layer of the water column and then decreasing sharply with depth (5 m depth; S1 Figs A and B). In Mejillones Bay, the chlorophyll-*a* contents varied between 0.03 and 3.5 mg/m^3^, recording the highest content at station E4BM (Fig 1; S1 Fig 1A) and the minimum surface content was observed at station E7BM (Fig 1; S1 Fig 1A). In Antofagasta Bay, the chlorophyll-a contents varied between 0.04 and 10.4 mg/m^3^, with the highest value occurring at station E7BA (Fig 1; S1 Fig 1B). The lowest surface content (1.1 mg/m^3^; 5 m depth) was observed at station E5BA (Fig 1; S1 Fig 1B).

At Mejillones Bay, the contents of NH_4_ ^+^ averaged 0.8 µM, with values that oscillated between 0.1 and 2.7 µM. Vertical profile patterns showed the minimum NH_4_ ^+^ content at a depth of 35 m at all stations (S2 Fig A). Similarly, the vertical profile concentrations of NO_2_ ^-^ showed a marked minimum at 35 m depth, whereas it had the highest content at 20 m depth (S2 Fig B). NO_2_ ^-^ averaged 0.5 µM at concentrations up to 2.8 µM. The NO_3_ ^-^ concentrations ranged between 0.1 and 23 µM, with the highest contents occurring at 35 m deep (S2 Fig C). Vertical profiles of PO_4_ ^3-^ showed a clear pattern of downward increment, reaching concentrations as high as 3.1 µM (S2 Fig D).

A more heterogeneous vertical and spatial distribution pattern of NH_4_ ^+^ was observed in Antofagasta Bay (S3 Fig A). Here, NH_4_ ^+^ concentrations averaged 0.4 µM, with concentrations that ranged between 0.1 and 1.5 µM (S3 Fig A). The NO_2_ ^-^ contents oscillated between 0.1 and 2.3 µM, averaging 0.3 µM (S3 Fig B). The NO_3_ ^-^ concentrations were lower at the surface (5 m depth; average 1 µM; S4 Fig C), whilst at deeper depths (35- 45 m depth), NO_3_ ^-^ contents reached up to 17 µM (S3 Fig C). The PO_4_ ^3-^ vertical profiles showed a pattern similar to that observed in Mejillones Bay at all stations, except for the vertical profile observed at station E1BA (S3 Fig D). At Antofagasta Bay, the PO_4_ ^3-^ contents varied between 0.5 and 3.7 µM and averaged 2.4 µM (S3 Fig D).

### 3.3. Chlorin Index

The chlorine index values in Mejillones and Antofagasta bays oscillated between 0.6 and 0.9 (S4 Fig A). In both bays, the chlorine index values showed a clear pattern of increase with depth, that is, the lowest values occurred in the shallower layer of the water column (5 m depth) and then increased progressively from 20 m to 45 m depth (S4 Figs A and B).

### 3.4. Total Organic Carbon

In Mejillones Bay, the total organic carbon (TOC) content ranged between 0.8 and 2 mg/L, resulting in an average content of 1.1 ± 0.4 mg/L (S4 Fig C; Table 1). The general vertical pattern distribution of TOC showed that the organic carbon content was higher in the first 20 m depth (S4 Fig C). At San Jorge Bay the average content of TOC was significantly higher than Mejillones bay, averaging 1.6 mg/L (Mann-Withney test; p = 0.001), with values oscillating between 0.7 and 3.7 mg/L (S4 Fig D).

**Table 1.** Mean values of δ^15^N-NO_3_ ^-^, δ^15^N of Phe and Thr, ΔTr and ΣV parameters, chlorin index, N*, AOU, and TOC obtained at the water columns of the Mejillones and Antofagasta bays. The data are presented as mean ± standard deviations.

| MEJILLONES BAY |  |  |  |  |  |  |  |  |  |
| --- | --- | --- | --- | --- | --- | --- | --- | --- | --- |
| Depth<br>(m) | $\delta^{15}\text{N}-\text{NO}_3^-$<br>(‰) | $\delta^{15}\text{N}-\text{Phe}$<br>(‰) | $\Delta\text{Tr}$ | $\Sigma\text{V}$ | $\delta^{15}\text{N}-\text{Thr}$<br>(‰) | Chlorine<br>Index | N* | AOU<br>($\mu\text{M}$) | TOC<br>(mg/L) |
| 5 | $12 \pm 0.4$ | $12 \pm 0.4$ | $0.9 \pm 0.1$ | $1.9 \pm 0.2$ | $15 \pm 1$ | $0.6 \pm 0.01$ | $-14 \pm 2$ | $22 \pm 15$ | $1.4 \pm 0.4$ |
| 20 | $13 \pm 0.2$ | $14 \pm 1.5$ | $1.9 \pm 0.3$ | $2.3 \pm 0.1$ | $13 \pm 2$ | $0.7 \pm 0.02$ | $-14 \pm 4$ | $161 \pm 50$ | $1.1 \pm 0.1$ |
| 35 | $15 \pm 0.1$ | $16 \pm 0.4$ | $2.1 \pm 0.3$ | $2.5 \pm 0.3$ | $10 \pm 3$ | $0.8 \pm 0.04$ | $-23 \pm 1$ | $230 \pm 18$ | $1.0 \pm 0.2$ |
| 45 | $16 \pm 0.2$ | $15 \pm 0.6$ | $1.3 \pm 0.5$ | $2.3 \pm 0.3$ | $8 \pm 2$ | $0.8 \pm 0.07$ | $-22 \pm 2$ | $242 \pm 8$ | $1.1 \pm 0.1$ |
| ANTOFAGASTA BAY |  |  |  |  |  |  |  |  |  |
| Depth<br>(m) | $\delta^{15}\text{N}-\text{NO}_3^-$<br>(‰) | $\delta^{15}\text{N}-\text{Phe}$<br>(‰) | $\Delta\text{Tr}$ | $\Sigma\text{V}$ | $\delta^{15}\text{N}-\text{Thr}$<br>(‰) | Chlorine<br>Index | N* | AOU | TOC<br>(mg/L) |
| 5 | $12 \pm 0.2$ | $12 \pm 0.4$ | $0.7 \pm 0.1$ | $1.6 \pm 0.1$ | $14 \pm 0.9$ | $0.6 \pm 0.01$ | $-9 \pm 3$ | $4 \pm 22$ | $2.1 \pm 0.6$ |
| 20 | $14 \pm 0.1$ | $15 \pm 1.6$ | $1.1 \pm 0.2$ | $2.0 \pm 0.2$ | $13 \pm 0.8$ | $0.8 \pm 0.07$ | $-23 \pm 2$ | $163 \pm 31$ | $1.6 \pm 0.7$ |
| 35 | $15 \pm 0.1$ | $16 \pm 0.4$ | $1.1 \pm 0.2$ | $2.0 \pm 0.2$ | $12 \pm 0.6$ | $0.8 \pm 0.05$ | $-32 \pm 3$ | $241 \pm 26$ | $1.6 \pm 0.4$ |
| 45 | $16 \pm 0.2$ | $15 \pm 0.6$ | $0.7 \pm 0.1$ | $1.8 \pm 0.1$ | $12 \pm 0.5$ | $0.9 \pm 0.04$ | $-34 \pm 3$ | $254 \pm 19$ | $1.7 \pm 0.4$ |

### 3.5. δ^15^N-AA patterns in suspended and sinking POM

In Mejillones Bay, the δ^15^N-AA values in sinking POM ranged between 5 ‰ to 27 ‰, averaging 17 ± 0.1 ‰ (Figs 3 A-C). Antofagasta Bay showed values that varied between 9 ‰ to 24 ‰, averaging 16 ± 0.1 ‰ (Figs 3 D-F). Non-significant differences in δ^15^N-AA values were found between both bays (Mann-Whitney test; p = 0.07). At both bays, trophic AA Ala, Val, Pro and Glu were the AA that showed the greatest ^15^N enrichment, (Figs 3 A-F). Leu and Ile were the trophic AA that exhibited the most depleted δ^15^N values (Figs 3 A-F). In both bays, the source AA, Gly, Thr, Phe, and Lys, averaged 14 ‰. In this group of AA, Phe and Lys showed the highest enrichment in δ^15^N values (Figs 3 A-C and 4 A-C).

**Fig 3.**
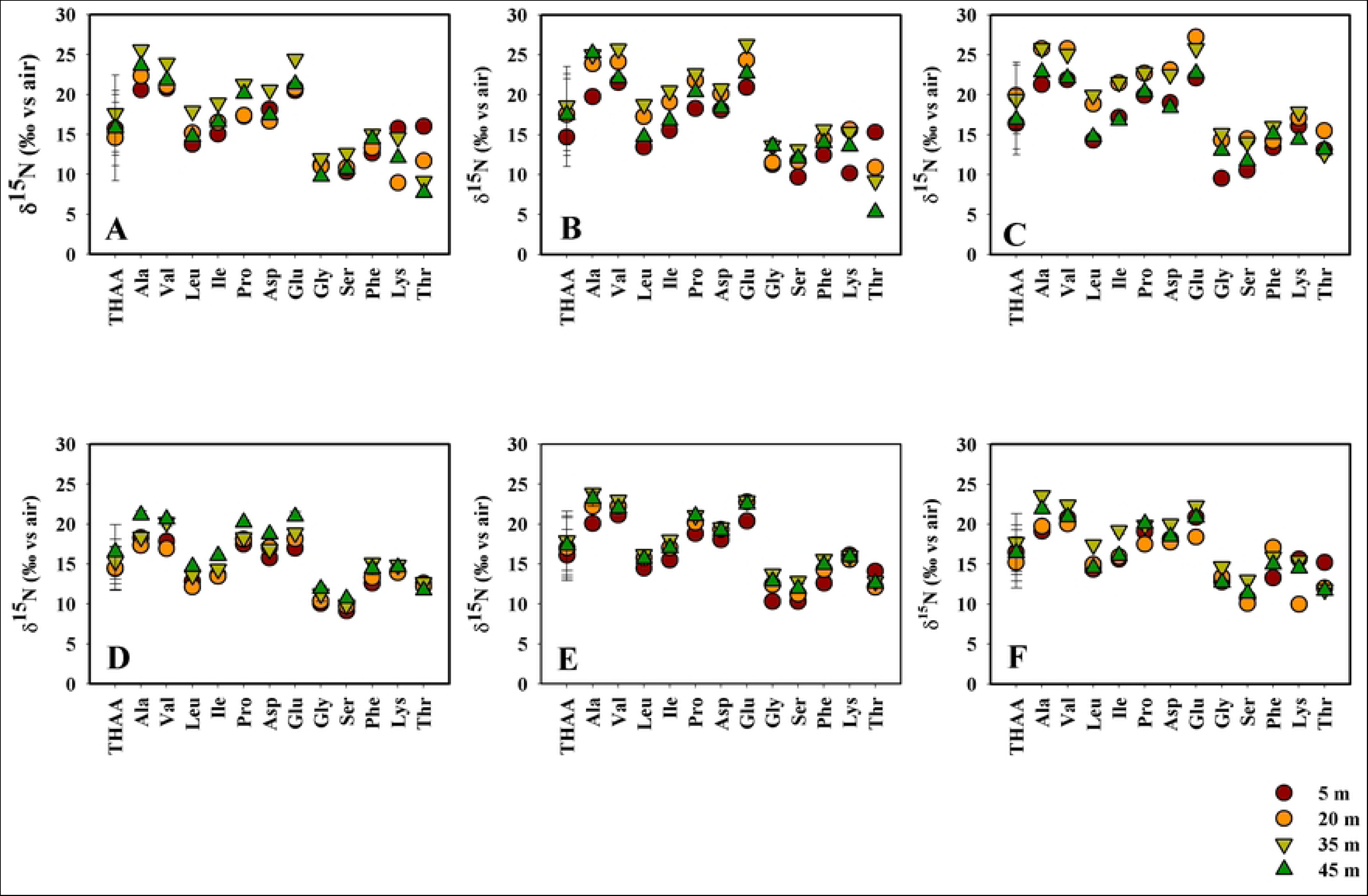
δ^15^N-AA signatures in suspended particulate organic matter in the water columns of (A-C) Mejillones Bay, and (D-F) Antofagasta Bay.

**Fig 4.**
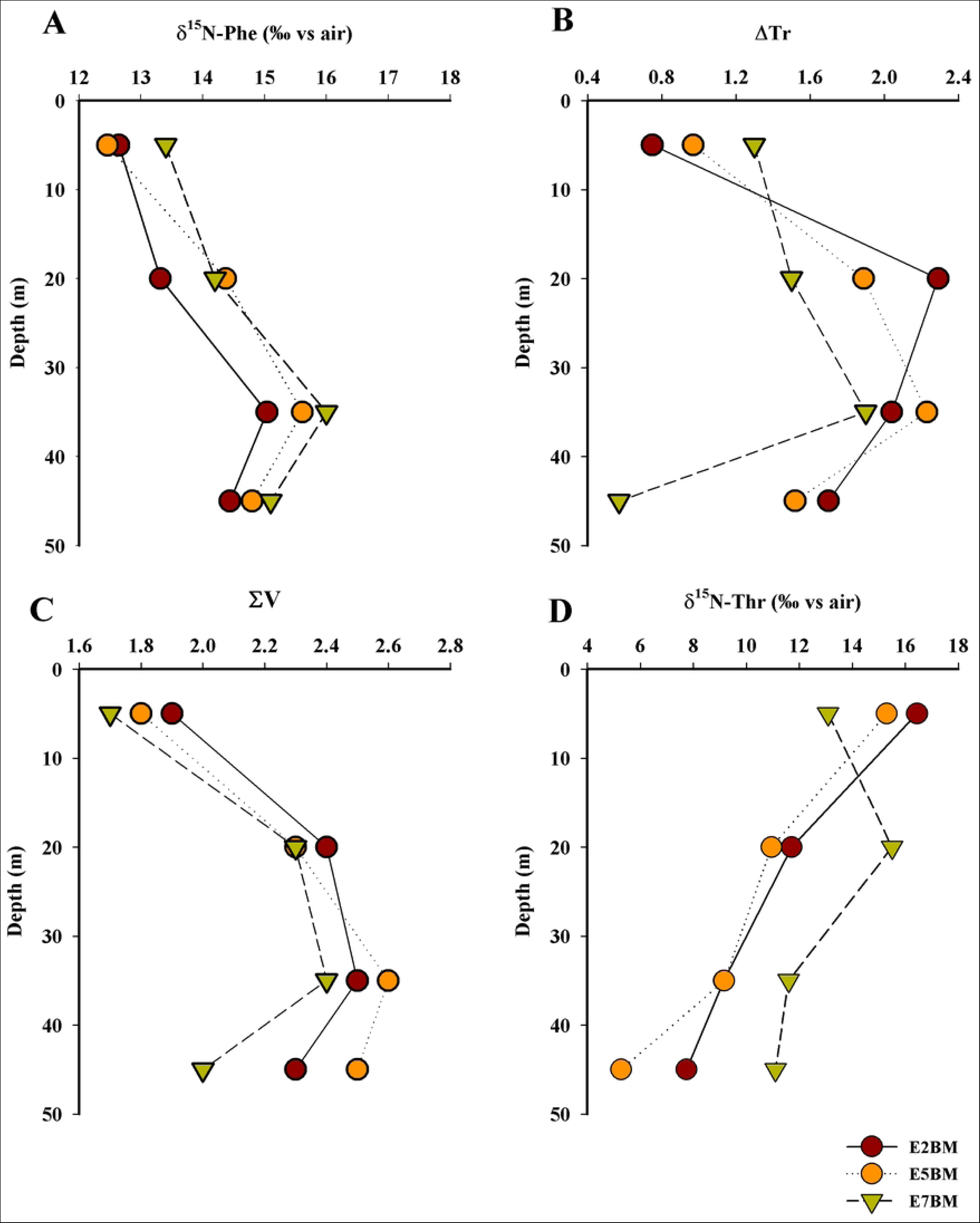
Vertical profiles of (A) δ^15^N Phe; (B) ΔTr, (C) ΣV; and (D) δ^15^N Thr in the water column of the Mejillones Bay.

In the northern sector of Mejillones Bay (E2BM; Fig 1), the δ^15^N of trophic AA values were highest at 35 m depth followed by the values found at 45 m depth, while the lowest values were evidenced generally at 5 m depth (Fig 3A). A similar pattern was observed in the central bay sector (E5BM; Fig. 3B). On contrary, at the southern sector of Mejillones Bay (E8BM; Fig 1), the highest δ^15^N of trophic AA values were observed at 20 and 35 m depth (Fig 3 C), while the lowest values of δ^15^N of trophic-AA occurred at 5 and 45 m depth (Fig 3C).

At the northern sector of Antofagasta Bay, the highest δ^15^N trophic-AA values occurred at 45 m depth (Fig 3D), whereas the lowest values of isotopic fractionation of Ala, Val, Leu and Ile were observed at 20 m depth and 5 m depth for Pro, Asp, and Glu (Fig 3D). In the central sector of Antofagasta Bay, the highest δ^15^N trophic-AA values occurred at both 35 and 45 m depths (Fig 3E), whereas the lowest δ^15^N trophic-AA values occurred at 5 m depth (Fig 3E). At the southernmost sector of the bay (E8BA; Fig 1), the highest δ^15^N trophic-AA values were found at 35 m depth (Fig 3F), whilst the lowest values were found between 5 and 20 m as well as at 45 m depth (Fig 3F).

Threonine (Thr) showed the opposite ^15^N isotopic behavior to the other trophic amino acids. At Mejillones Bay, Thr showed higher values of δ^15^N at surface, while, in the deeper layers, this amino acid was more depleted (Figs 3A-F). At north of Mejillones bay (E2BM; Fig 1), the δ^15^N isotopic abundance of Thr was higher at surface water (16 ‰; 5 m depth; Fig 3A), whilst at 20 m depth this amino acid was more depleted (δ^15^N = 11 ‰). At 35 and 45 m depth, the δ^15^N values for Thr were 9 ‰ and 8 ‰, respectively (Fig 3A). At the E5M station (center of the bay; Fig. 1), Thr was more depleted in δ ^15^N at 45 m depth (δ^15^N = 5 ‰; Fig 3A), and more enriched at 5 m depth (15 ‰; Fig 3B). At the southernmost station (E7BM; Fig. 1), Thr was enriched up to a depth of 20 m (Fig 3C). At Antofagasta Bay, the δ ^15^N values of Thr, were similar at the stations E2A and E5A, without a clear pattern of decrease or increase in δ^15^N -Thr values with depth, showing values that varied between 11 ‰ and 14 ‰ (Figs 3 D and E). At station E8BA, Thr was δ ^15^N enriched at surface water (15 ‰; 5 m depth; Fig 3C).

The sinking POM collected from the sediment traps (45 m depth), showed values and isotopic fractionation patterns similar to those observed in the deeper layers of the water columns in both bays (Table 1). In Mejillones Bay, the average δ^15^N -AA value was 16 ± 2. Trophic AA were the most enriched in, with an averaged δ^15^N value of 22 ± 3 ‰ (Table 1). The average δ^15^N value of the source AA was 15 ± 1 ‰. In Antofagasta Bay, the averaged δ^15^N -AA value was 18 ± 3 ‰. Trophic AA δ^15^N values averaged 21 ± 2‰, while source AA averaged 14 ± 1‰ (Table 1). As well as in the suspended particulate material, the δ^15^N- Thr values were more depleted, compared to the trophic and source AA in both bays. As well as in the suspended particulate material, the averaged δ^15^N of sinking Thr were more depleted, compared to the trophic and source AA in both bays (Table 1)

### 3.6. ΔTr and ΣV in suspended and sinking POM

At Mejillones Bay, the ΔTr values varied between 0.6 and 2.3, whereas at Antofagasta Bay the values were significantly lower than the observed at Mejillones Bay, with values that oscillated between 0.5 and 1.3 (Mann-Whitney test, p = 0.01) (Figs 4C and 5C). The vertical profile patterns were similar in both bays, showing the highest ΔTr values between 20 and 35 m depth (Figs 4C and 5C), except for the observed at station E2BA, where the increase in ΔTr values occurred deepest (e.g., 35 m depth; Figs 1A and 5C). No significant differences in ΔTr values were observed between stations both in Mejillones Bay and neither in Antofagasta Bay (Wilcoxon Test, p > 0.05).

**Fig 5.**
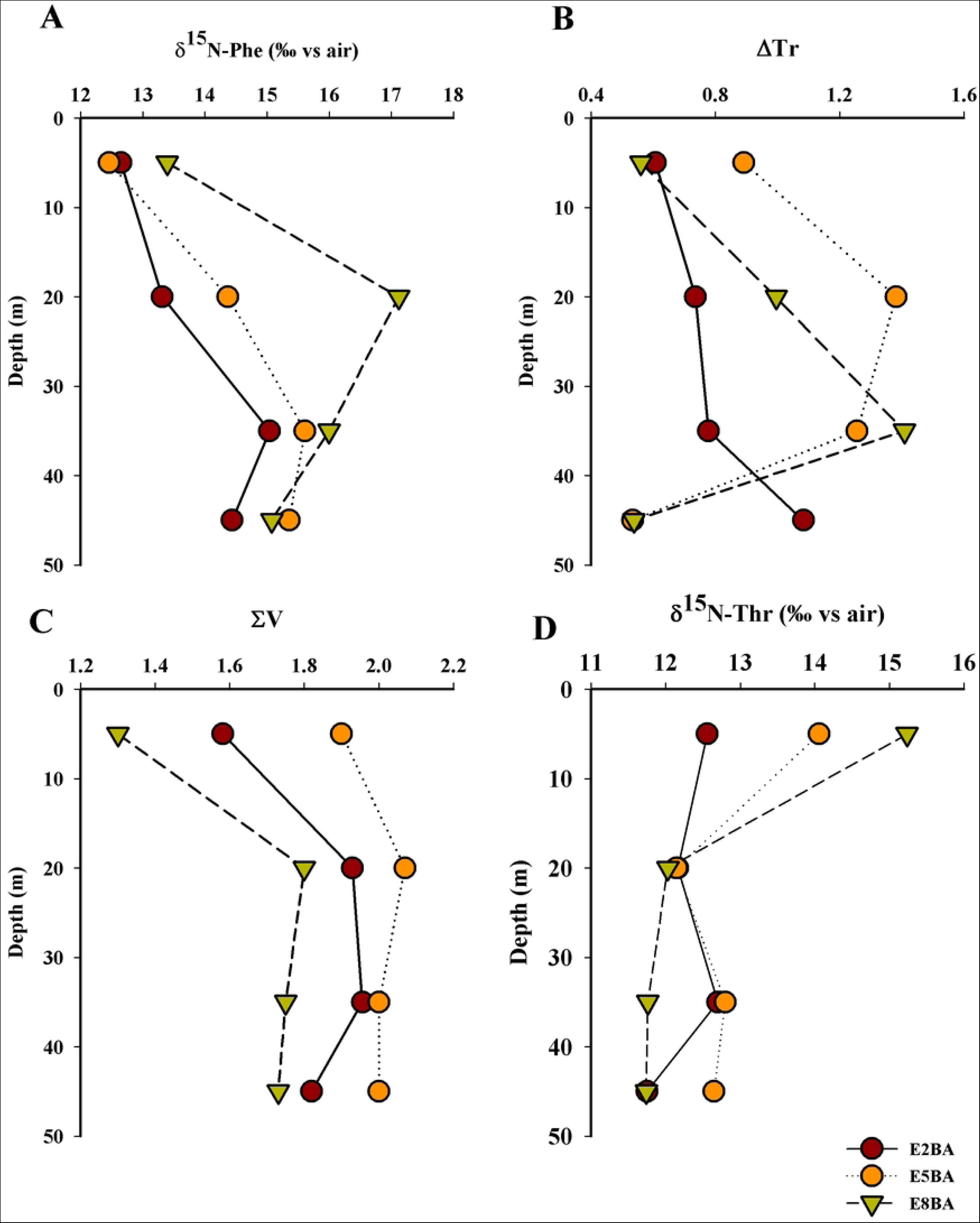
Vertical profiles of (A) δ^15^N-Phe; (B) ΔTr, (C) Σ*V*; and (D) δ^15^N Thr in the water column of the Antofagasta Bay.

At Mejillones Bay, the ΣV values averaged 2.3 ± 0.05 (Table 2). In the northern sector of the bay (station E2BM; Figs 1 and 4D), the ΣV values ranged between 2.3 and 2.7, showing a clear increment pattern with depth (Fig 4D). At station E5BM, the ΣV values increased continuously from 2.1 in the shallower water layer to 2.5 at 35 m depth (Fig 4D), then decreased slightly to 2.3 at 45 m depth (Fig 5D). A different vertical pattern was observed at station E7BM, located in the southernmost sector of the bay (Fig. 1). A subtle increase in ΣV values was observed from 5 to 20 m depth (2.2 to 2.3; Fig 5D), followed by a sharp decrease to 1.9 at 35 m depth (Fig 5D). Significant differences were found between the ΣV values at stations E2BM and E7BM (Wilcoxon Test, p = 0.04).

**Table 2.** Mean values of δ^15^N-AA, ΔTr and ΣV parameters, and downward fluxes values obtained from sediment traps deployed at Mejillones and Antofagasta bays.

| MEJILLONES BAY |  |  |  |  |  |  |  |  |  |  |  |  |  |  |  |
| --- | --- | --- | --- | --- | --- | --- | --- | --- | --- | --- | --- | --- | --- | --- | --- |
| $\delta^{15}\text{N}$-AA (‰) | | | | | | | | | | | | | Parameter | | |
| THAA | Ala | Val | Leu | Ile | Pro | Asp | Glu | Gly | Ser | Phe | Lys | Thr | $\Delta\text{Tr}$ | $\Sigma V$ | Downward flux<br>($\text{mg}/\text{m}^2/\text{day}$) |
| 16 | 25 | 26 | 18 | 21 | 22 | 20 | 26 | 12 | 13 | 16 | 19 | 8 | 2.1 | 2.4 | $9 \pm 2$ |
| ANTOFAGASTA BAY |  |  |  |  |  |  |  |  |  |  |  |  |  |  |  |
| $\delta^{15}\text{N}$-AA | | | | | | | | | | | | | Parameter | | |
| THAA | Ala | Val | Leu | Ile | Pro | Asp | Glu | Gly | Ser | Phe | Lys | Thr | $\Delta\text{Tr}$ | $\Sigma V$ | Downward flux<br>($\text{mg}/\text{m}^2/\text{day}$) |
| 15 | 24 | 22 | 16 | 18 | 22 | 20 | 23 | 13 | 13 | 15 | 16 | 11 | 1.2 | 1.9 | $11 \text{g} \pm 1$ |

In Antofagasta Bay, significantly lower ΣV values were observed (Mann-Whitney test, p < 0.01), with an average of 2.1 (Table 2). The highest ΣV values were recorded at the center of the bay (station E5BA; Figs 1 and 5D). These variations in ΣV values between stations and bays suggest differences in the intensity of microbial processing and trophic transfer of POM.

### 3.8. Natural abundance of δ^15^N-NO_3_ ^-^ isotopes

At Mejillones and Antofagasta bays, the δ^15^N abundances values of dissolved NO_3_ ^-^ ranged between 11 ‰ and 14 ‰ (Table 2). No significant differences were observed between the two bays (Mann-Whitney test; p = 0.6; Table 1). A clear enrichment trend with depth was observed at both bays, with an average increase of 2.6 ± 0.01 ‰. The lowest values occurred at the surface (average δ^15^N-NO_3_ ^-^ = 12 ± 0.1 ‰ at 5 m depth), while the highest values were observed at 45 m depth (δ^15^N-NO_3_ ^-^ = 15 ± 0.01 ‰) (Table 1).

### 3.9. Downward particulate organic carbon flux

In Mejillones Bay, after 30 days of deployment, the sediment traps located at the center of the bay (E5BM; Fig 1) collected 97 mg/L of particulate organic carbon (POC), while the collectors deployed at the southernmost station (E7BM; Fig 1) retained 135 mg/L of POC. In Antofagasta Bay, the highest content of POC collected in the traps occurred at the center of the bay (E5BA; Fig 1), with a POC concentration of 156 mg/L, followed by 120 mg/L in the sediment trap located at the southernmost station of the bay (E8BA; Fig 1). Based on the aspect ratio of the trap array and the deployment time, a mean downward flux value of 9 ± 2 mg/m²/day of POC was estimated for Mejillones Bay (Table 2). In Antofagasta Bay, the estimated mean flux of POC was 11 ± 1 mg/m²/day (Table 2).

## 4. DISCUSSION

### 4.1. Source and origin of POM in Mejillones and San Jorge bays

The enriched δ^15^N-NO_3_^-^ values observed in the waters of the Mejillones and Antofagasta bays (Table 1), exceed the average δ^15^N-NO_3_^-^ value obtained in Pacific oceanic waters (e.g., 5‰), but are similar to the values of δ^15^N-NO_3_^-^ previously reported in coastal waters influenced by OMZ waters of northern and central southern of Chile (56,57), and to those reported from other marine oxygen-depleted waters of the open ocean (58–60). The fractionation of nitrogen isotopes associated with denitrification leads to significant enrichment in the ^15^N of residual NO_3_ ^-^ (17,58,59,61), generating a useful isotopic indicator of denitrification. A similar isotopic discrimination occurs with phytoplankton, which preferentially absorbs the isotopically lighter NO_3_ ^-^ containing ^14^N (58, 59), leaving the remaining dissolved inorganic nitrogen more enriched in ^15^N as a result of phytoplankton uptake (64). Therefore, we hypothesize that the δ^15^N-NO_3_ ^-^ values found in both waters columns were modulated mainly by biological mechanisms involving NO_3_ ^-^ assimilation processes and dissimilative NO_3_ ^-^ reduction (56,57). The surface layers of both bays were completely oxygenated (Figs 2 C and H), hosting a conspicuous zone of a minimum NO_3_ ^-^ content (S2 Fig C and S3 Fig C), pointing out to photosynthetic NO_3_ ^-^ assimilation as the potential process responsible for the observed δ^15^N-NO_3_ ^-^ and δ^15^N-Phe signals at surface oxic waters (Figs 3 A-F). Therefore, and according to our data, it appears that the surface δ^15^N-NO_3_ ^-^ is not only modulated by the δ^15^N-NO_3_ ^-^ of the upwelled NO_3_ ^-^ but also by the absorption and assimilation of NO_3_ ^-^ (56).

Another alternative explanation for the most depleted δ^15^N-NO_3_ ^-^ values observed at the surface (12 ‰; Table 2), compared with the deeper δ^15^N-NO_3_ ^-^ values, could be the addition of isotopically lighter NO_3_ ^-^, originated by the oxidation of recently fixed N (65). Nitrogen fixation has been previously reported in the waters of the oxygen minimum zone of central-southern Chile (66). This process has the potential to account for approximately 20% of new N production in surface waters associated with oxygen minimum zones (67). Despite that the surface δ^15^N-NO_3_ ^-^ values were lower than those observed at the deeper layers of both bays, these were more enriched than the values observed in similar depths of the waters of the Mediterranean Sea and North Atlantic (60,68) and the average value for deep NO_3_ ^-^ in the world ocean (i.e., 5 ‰) (59,64). The narrow vertical segregation of more depleted values of δ^15^N-NO_3_ ^-^ and minimum contents of NO_3_ ^-^ observed at the surface could be explained by the intense (strong and thin) thermoclines characteristic of upwelling shadows bays (28,33,69), and upwelling traps bays (29,38), often supporting a thin-layer subsurface of chlorophyll maxima (70–72), as was observed in the study area (S1 Figs A and B; S2 Fig C and S3 Fig C).

On the contrary, the observed enrichment of δ^15^N-NO_3_ ^-^ with depth, alongside the progressive vertical depletion of oxygen content in both bays (Figs 2C and H; Table 1), appears to have been driven by microaerophilic and/or anaerobic metabolic processes. These processes are associated with N loss and are primarily carried out by microbes due to the expected scarcity or absence of metazoans in hypoxic/suboxic/anoxic waters (73–76). These conditions were achieved in the deeper water layers, where the strongest oxygen depletion and more negative N* values were recorded (Figs 2 C and H; Table 1). Therefore, NO_3_ ^-^ was microbially used as the thermodynamically most favorable terminal electron acceptor during the degradation of the organic matter, resulting in the observed isotopically heavier and deeper residual NO_3_ ^-^, as has been reported in other marine settings (54,63,72– 74; 75).

In addition to biological forcings, physical and hydrodynamic forcings could also have contributed to and modulated the observed vertical pattern distribution of δ^15^N-NO_3_ ^-^ in the water columns of both bays. The T-S diagrams show the predominance of warmer, more saline, and more oxygenated Surface Subtropical Water (SSTW) and its modified form, as well as modified Equatorial Subsurface Water (ESSW), within both bays (Figs 2E and J). The occurrence of SSTW and AESS in Mejillones and Antofagasta bays has been previously demonstrated, with vertical distributions of mixing percentages of water masses similar to those observed in this work (40,80–82).

The mesoscale offshore circulation pattern of the currents in front of both bays is one of the mechanisms responsible for the distribution of water masses and their mixing, given that the Mejillones Peninsula constitutes a transition zone where surface water masses of subtropical and sub-Antarctic origin converge (83). As a result of this convergence, the SSTW would preferentially enter in summer, given that this mass of water stands out in the warm season (40). However, the presence of AESS in the deeper water layers of both bays can be attributed to the upwelling mechanism (29,30,33,34,37,84). The hydrodynamic interaction between these waters and their consequent mixing could have generated a dilution effect in the δ^15^N-NO_3_ ^-^ signal towards the surface, which may work to further homogenize the δ^15^N of the upwelled NO_3_ ^-^. Consequently, one might posit that, as the upwelled waters move away from the core of the OMZ, to more oxygenated waters, the oxygen would progressively increase and the δ^15^N-NO_3_ ^-^ would decrease, likely due to the mixing between the deeper, oxygen-depleted and high δ^15^N-NO_3_ ^-^ upwelled waters (ESSW) and low δ^15^N-NO_3_ ^-^ and more oxygenated SSTW, in a similar hydrodynamic mechanism to the previously proposed by De Pol-Holz et al. (79).

The δ^15^N-NO_3_ ^-^ values observed at the first 20 m depth of the water columns of both bays (mean δ^15^N-NO_3_ ^-^ value = 12 ± 0.4 ‰; Table 1) were similar to the surface δ^15^N-Phe-POM values, (Table 1), which suggests strongly that POM has an autochthonous photosynthetic origin, that is fueled mainly by nitrate enriched in ^15^N, upwelled from the deeper oxygen-depleted waters, so we infer that POM formed *in situ* dominated the POM pool of the study area. In these waters, primary productivity is fundamentally restricted to the first 20 meters of the water column (32), which is conceptually and operationally consistent with the vertical distribution patterns of chlorophyll-*a* contents and δ^15^N isotopic values of Phe and NO_3_ ^-^ observed in the water columns of both bays (S1 Figs A and B; Table 1).

The potential contributions of organic matter of anthropogenic origin cannot be ruled out given the intense anthropogenic pressure to which both bays are subjected, which is related to mining industry port operations, desalination plants, artisanal fisheries, artificial touristic beaches, and sewage discharge (85–87). The δ^15^N-THAA values of POM found in the study area, considered here as bulk δ^15^N-POM (see methods section for the operational definition), oscillated between 15 ± 3 ‰ to 17 ± 2 ‰ (Figs 3 A-F). These values enriched in ^15^N in the suspended and sinking POM are expected for waters influenced by oxygen minimum zones (60,79,88), but are also similar to the values observed in wastes that have experienced anthropogenic disturbances, which are isotopically rich in heavy nitrogenous components, such as N derived from human wastewater and livestock (e.g., δ^15^N of 10‰ to 22‰) (89,90), which makes it difficult to discriminate between anthropogenic and natural sources using only our bulk δ^15^N -POM data. However, our auxiliary bulk δ^13^C-POM data were less ambiguous, yielding average values of −22 ± 6 ‰ and −21 ± 5 ‰ for Mejillones and Antofagasta bays, respectively (S5 Fig), which are commonly assigned to organic matter of marine origin in mid- and low-latitude regions (91–93). This reinforces our inferences, based on our δ^15^N-NO_3_ ^-^ and δ^15^N-Phe data, about the origin and sources of POM in the study area, and confirms the dominance of POM of marine origin in the waters of both bays.

Contrarily, it highlights the fact that, in both bays, the δ^15^N-Phe values in the POM increased with depth, more than the ^15^N-NO_3_ ^-^ values, mainly between 20 to 35 m depth (i.e., the base of the oxycline) (Figs 4A and 5A; Table 1). Although it has been widely demonstrated that δ^15^N-Phe values exhibit little fractionation in experimental cultures with fungi, bacteria and archaea using organic substrates (13,94,95), it has also been observed that δ^15^N-Phe values can change when peptide bonds are broken during microbial degradation of organic matter (23). The observed synchronic δ^15^N enrichment of trophic AA and Phe with depth (Figs 3A-F, 4A, and 5A), is similar to that observed by Hannides et al. (22), who suggested that this AA δ^15^N isotopic enrichment pattern is the result of the degradation of suspended organic matter through external hydrolysis (Rayleigh fractionation), so that the δ^15^N values of trophic and source AA (as well as all others AA), increase uniformly with external hydrolytic enzymatic activity. In the oxic and suboxic waters of northern Chile, microbial peptide hydrolysis occurs at rates similar to or faster than amino acid uptake in the water column (96,97). Therefore, the evidence of synchronic isotopic enrichment of ^15^N-Phe and trophic AA could be due to intense extracellular enzyme activity on photosynthetic suspended and sinking POM through the oxycline in both bays. Extracellular enzyme activity has previously been observed in various marine particles, including suspended particulate matter (98,99), marine snow (100,101), and sinking particles collected in sediment traps (102,103).

### 4.2. Vertical and spatial patterns of trophic transfer and recycling of POM in Mejillones and San Jorge bays

Mejillones and Antofagasta bays, two semi-enclosed bodies of water located on the Mejillones Peninsula, are strongly influenced by mesoscale offshore processes, which are modulated by the intensity and direction of atmospheric circulation. However, these bays differ significantly in their bottom topography and geographic orientation. These differences result in contrasting circulation patterns, stratification, residence times, and hydrographic processes related to upwelling (29,30,33,35,36). Moreover, these bays experience distinct and varying degrees and types of anthropogenic pressure along their coastlines (87).

In this context, we evaluated whether these contrasting oceanographic and topographic conditions could produce differences in trophic transfer and the degree of heterotrophic reworking of POM. Our ^15^N-AA isotope-based data revealed that ΔTr and ΣV parameter values resemble the expected vertical distribution of photosynthetic and microbial activity for these waters, modulated by the vertical physical and chemical gradients found in the upwelling ecosystem of Chilean coastal waters (57,97,104–111).

At the surface layers of both bays (5 m depth), the averaged ΔTr and ΣV values were less than 0.9 and 1.9, respectively (Table 1), indicating the dominance of a phytoplanktonic origin for the POM and zooplanktonic processing of the detrital organic material (1). This observation is of biogeochemical and ecological significance, given that zooplankton grazing is one of the key processes by which phytoplankton carbon is transferred to higher trophic levels in marine ecosystems (108,109, 110,111). Thus, the ΔTr values suggest that the transfer of POM from phytoplankton to higher trophic levels of the pelagic food web is predominantly mediated by zooplanktonic grazing in the shallowest water layers of Mejillones and Antofagasta bays. This finding aligns with a previous report by Valdés et al. (86), which showed that in highly productive waters, phytoplanktonic carbon is efficiently transferred through grazing by copepods (116). According to the vertical profiles of chlorophyll and ΔTr, zooplanktonic grazing could be segregated in the well-oxygenated and food-rich photic layer (surface layer), due to the intrusion and occurrence of deeper oxygen-depleted waters resulting from coastal upwelling in the coastal waters of northern Chile (28,33,84). The vertical distribution of zooplankton, delimited by dissolved oxygen availability, has been previously observed in the northern upwelling region of Chile (117) and the oxygen minimum zone of the Eastern Tropical North Pacific (118).

On the contrary, in both bays, the δ^15^N values of trophic and source AA were more depleted at the surface (i.e., 5 m depth) compared to the values observed below 20 m depth (Figs 3 A-F). These observations align with earlier studies that documented a substantial increase in the ^15^N content of suspended particles with depth in both marine and lacustrine water columns (1,12,17,22). This enrichment of the heavier N isotopes is explained by the degradation of labile ^15^N-depleted organic material, leaving behind a ^15^N-enriched pool of suspended material (22). This interpretation suggests that heterotrophic resynthesis and trophic transfers are the main mechanisms driving the ^15^N enrichment of AA in suspended and sinking POM in the water columns of Mejillones and Antofagasta bays (Figs 4 A-D and 5 A-D).

This conclusion is further supported by the vertical distribution of ΔTr and ΣV parameter values (Figs 4C and 5C). In the water columns of both bays, trophic transfer and POM resynthesis increased with depth, reaching their highest values at the base of the oxycline (i.e., bellow 35 m depth) (Figs 4B and 5B). Additionally, the increases in NH_4_ ^+^ and PO_4_ ^3-^ contents, enhanced apparent oxygen utilization, and decreased pH and TOC content with depth (Figs 2 E and I; S2 Figs A and D; S3 Figs A and D; S5 Fig and S6 Fig D; Table 1) support the notion of more intense heterotrophic respiration of POM at the base of the oxycline, where dissolved oxygen depletion is most severe. These observations align with to the reported by Gonzalez et al. (119), who found that microbial community inhabiting oxygen depleted waters of Chile has the same or even greater potential heterotrophic metabolic activity than the microbes thriving in the more oxygenated water layers. This is consistent with previous observations in the coastal waters of northern Chile, where temporal and vertical variability in levels of N, P, and Si, as well as N and Si ratios, indicate significant dynamics in the recycling processes of organic matter in the water column (31,82).

These findings strongly demonstrate the existence of an active heterotrophic microbial assemblage, with activity appearing to be more intense at through the oxyclines in both bays (e.g., bellow 20 m depth). Recently, Srain et al. (111) demonstrated through incubation experiments and field measurements the presence of a complex heterotrophic microbial consortium in the suboxic/anoxic coastal waters of the upwelling ecosystem of central Chile. In both bays (approximately 20–45 m depth; Fig 1), the dissolved inorganic N anomalies (NO_3_ ^-^ + NO_2_ ^-^+ NH_4_ ^+^; N*) (Table 1) were likely associated with microaerophilic and anaerobic metabolism in the water column N cycle. The N* values indicated an NO_3_ ^-^ deficit throughout both oxyclines, with similar negative N* values previously recorded in the upwelling system located between 21° and 33°S, attributed to the occurrence of denitrification and anammox metabolism (79,120,121).

At the deeper water layers of both bays (i.e., 45 m depth), the δ^15^N-Phe values were more depleted compared to the δ^15^N-Phe signal of POM obtained from the shallower water layers (i.e., 2 ± 1 ‰; Figs. 4A and 5A; Table 1). These deeper values were closer to the ambient δ^15^-NO_3_ ^-^ values and coincided with more negative N* values and decreased ΔTr and ΣV parameters, a pattern similar to that found in autotrophic biomass (Table 1). A similar pattern was observed in chemoautotrophic bacteria grown on an NH_4_ ^+^ substrate, which showed phytoplankton-like ΔTr and ΣV values (13). This suggests a potential occurrence of chemoautotrophic metabolism. Chemoautotrophic processes, such as annamox and HS^-^ oxidation coupled with NO_3_ ^-^ reduction, have been observed in the oxygen-depleted waters of northern and central-southern Chile (122–124).

Our δ^15^N-AA derived data also revealed significant differences in the spatial distribution and intensity of these processes between and within both bays, indicating a heterogeneous spatial pattern. Specifically, Mejillones Bay exhibited significantly higher values of ΔTr and ΣV compared to Antofagasta Bay (Mann-Whitney test; p = 0.01), implying substantial differences in the intensity of respiration of suspended detrital organic material, trophic transfer of organic carbon, and the degree of degradation, amount, and quality of the exported organic material (Tables 1 and 2). For example, the vertical profiles of δ^15^N-Phe demonstrated that the vertical extension of enzymatic hydrolytic activity in Mejillones Bay occurred over a broader water layer (approximately 15 m wide; Fig. 4A) compared to Antofagasta Bay. This broader enzymatic activity coincided with a lower export flux of POM in Mejillones Bay (Table 2). The increase and attenuation of the vertical flux of POM observed in both bays could have been influenced as well by differences in the occurrence and intensity of abiotic processes such as mineral dissolution, biotic processes mediated by enzymatic kinetic associated with microbial reactions (125–127) and the intensity of zooplankton grazing (128), as has been observed previously in mesopelagic water columns (129). This is consistent with the lower values of ΔTr and ΣV, and the higher fluxes of exported POM collected from the sediment traps deployed at Antofagasta Bay.

These findings reveal a lower trophic transfer status and a lesser degree of heterotrophic reworking of POM, which could result in a higher downward flux of POM at Antofagasta Bay compared to Mejillones Bay (Table 2). This inference is further supported by the lower δ^15^N-Thr depletion below the 20 m depth observed in Antofagasta Bay compared to Mejillones Bay (Mann-Whitney test; p = 0.01).This potentially indicates a lower metabolic reworking of POM due to a lower degree of trophic transfer, as Thr ^15^N isotope fractionation behaves differently from other AA, often showing significant depletion during trophic transfer (130). Specifically, Antofagasta Bay exhibited an average offset between surface and bottom δ^15^N-Thr values of 2 ‰, whereas in Mejillones Bay, this offset reached 8 ‰ (Figs 4D and 5D).

This heterogeneity in the spatial distribution of the POM processing and trophic transfer described above was also observed within both bays (Figs 4A-D and 5A-D). For instance, in Mejillones Bay was evident a clear spatial pattern of decrease in the values of ΔTr, ΣV and δ^15^N-Thr in a north-south direction, while in Antofagasta Bay, this pattern was not so clear, although some spatial differences were observed (Figs 4 B-D and 5 B-D).

The similitude between the vertical physical-chemical structure of the water column of both bays (Figs. 2 A-J; Mann-Whitney test; p > 0.05; S1 table), in addition to the lack of significant associations between the ΔTr and ΣV, with temperature, salinity, dissolved oxygen and pH (Spearman r; p > 0.05; S6 Fig), suggested to us that other unexamined factors may be responsible for the spatial heterogeneity in trophic transfer and heterotrophic recycling of POM described earlier.

Potentially, local-scale circulation patterns, residence times, stratification, and the occurrence of thermal fronts, in addition to differences in coastal geometry and bottom topography, could have influenced the contrasting conditions and spatial variability in trophic transfer levels and POM recycling between both bays. For instance, the geographical orientation and coastline topography of Mejillones Bay can create anomalously high-temperature zones, generating thermal fronts such as upwelling shadows (131), which enhance water residence times and reduce horizontal transport within the bay (33). Conversely, Antofagasta Bay is classified as an upwelling trap bay, where winds blowing into the bay from upwind of a prominent headland often trap water due to its full exposure to the alongshore shelf jet (28–30,33). When comparing both bays, it was observed that Mejillones Bay presented on average a lower surface temperature, a lower dissolved oxygen inventory and a lower stratification of the water column than Antofagasta Bay (Figs 6 A-C). In Mejillones Bay, the surface temperature was colder at the head of the bay, while the warmest temperatures were evident in the center of it (Fig 6A). A similar spatial pattern was observed for the dissolved oxygen inventory, highlighting the head area of the bay as the most oxygen-depleted sector (Fig 6B). Greater stratification was also observed from the center towards the mouth of the bay (Fig 6C). On the contrary, it was observed that in Antofagasta Bay, the surface temperature was higher at the head of the bay (Fig 6A), with a less clear spatial pattern in terms of dissolved oxygen inventories, highlighting a maximum integrated content in the center of the bay (Fig 6B), and a band of greater stratification on the outer face of the bay (Fig 6C). Previously, it has been shown that these physical and hydrodynamic forcings, induced by the orientation, geometry, size and general geomorphology of these coastal formations determine and modulate biological processes such as primary productivity, community respiration (107, 130, 131–137), retention and recruitment of planktonic propagules as well as explain patterns in pelagic and benthic habitats in the upwelling bays (36,131,139,140). Here, we suggest that these forcings also have the potential to trigger contrasting spatial patterns, at local scale, in the intensity of heterotrophic reworking and trophic dynamics of suspended and sinking POM. However, these observations are speculative and warrant more specific complementary hydrodynamic studies at local scale to fully address these hypotheses, which remain open. Our results challenge the paradigm that the vertical physical-chemical structure of the water column alone ultimately modulate the spatial patterns and intensities of organic carbon cycling in the coastal pelagic environments, and highlight the interactively influence of physical, chemical, and biological dynamics alongside with the heterogeneous nature of topographic morphology and geographic orientation of the upwelling bays systems.

**Fig 6.**
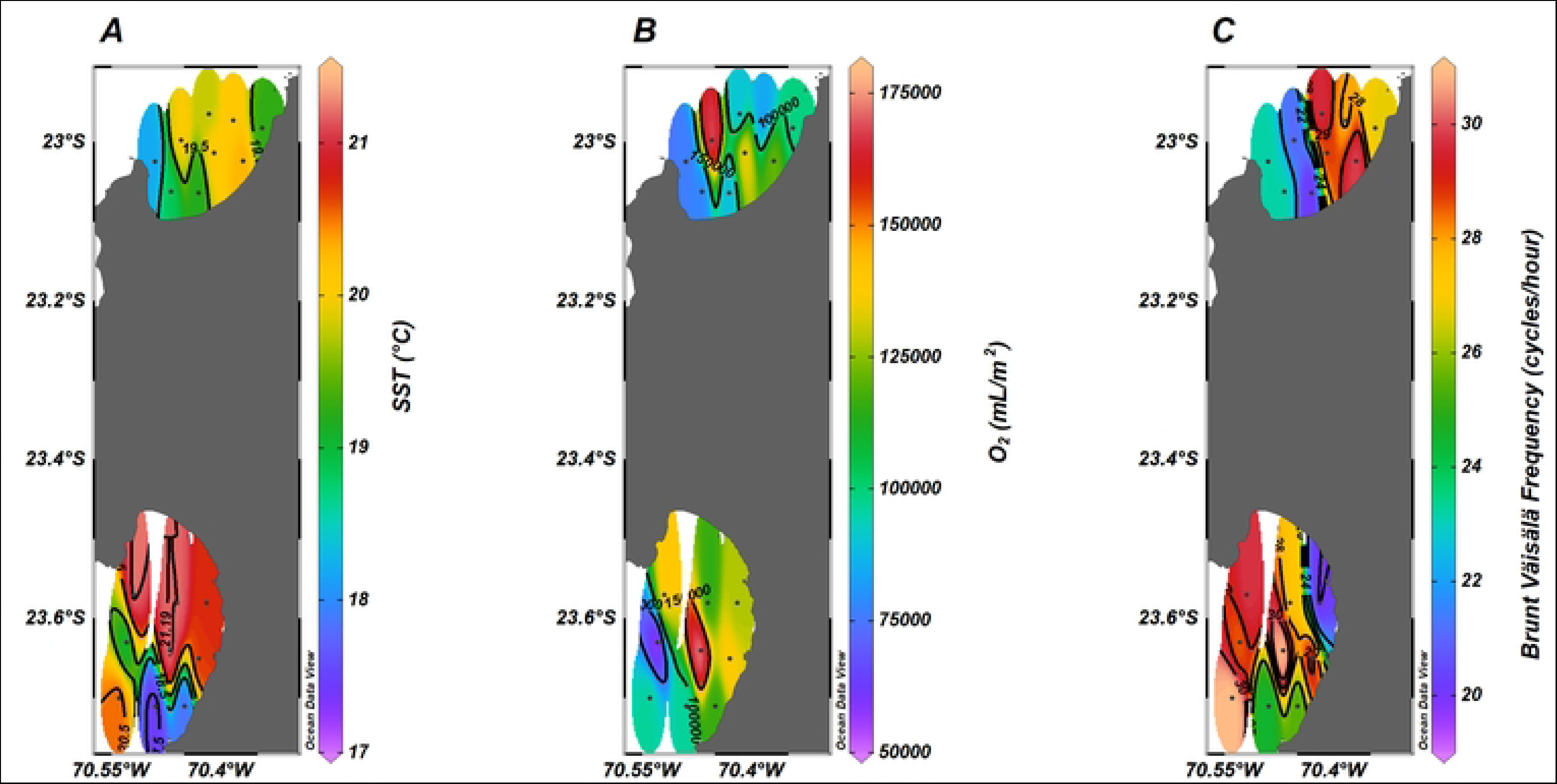
Spatial distribution of (A) sea surface temperature (SST); (B) dissolved oxygen inventories and (*C) stratification of the water column in the bays of Mejillones and Antofagasta. (*) The reported values of the Brunt-Väisälä Frequencies correspond to the value averaged in the first 20 meters of the water column.

## 5. CONCLUSIONS

The observed isotopic signatures of δ^15^N-AA patterns, serve as robust indicators for investigating the sources and extent of heterotrophic processing of POM in highly productive coastal environments. Variations in δ^15^N values of individual AA, reflective of trophic transfers, and heterotrophic reworking persist in suspended and sinking POM, alongside additional alterations that suggest subsequent microbial transformations post-incorporation into particles. The δ^15^N -Phe and δ^15^N-NO_3_ ^-^ values, confirms the photosynthetic origin of POM, predominantly derived from autochthonous sources. This organic material is synthesized in the uppermost water column (above 20 m depth) and undergoes heterotrophic processing primarily through the oxycline (approximately 20–35 m depth). Analysis of ΔT and ΣV parameters reveals significant spatial variability, at local scale, in trophic transfer and microbial heterotrophic reworking of POM. These findings underscore the heterogeneous nature of organic matter transfer and recycling processes within these upwelling bay systems. Such insights emphasize the necessity to consider each bay as a distinct functional oceanographic unit when formulating environmental policies and management strategies.

## ACKNOWLEDGMENTS

This research was funded by “Asociación de Industriales de Mejillones”, Chile. We acknowledge the support provided by professors Juan Ávila Donoso and Pedro Cortés Peña of the Chemistry, Bioinorganic and Analytical Laboratory of the Faculty of Basic Sciences of the University of Antofagasta. We are grateful to the crew of the L/C ANAGO for help during sampling, and the Marine Ecologist Maritza Malebrán for her technical support.

## SUPPORTING INFORMATION CAPTIONS

**S1 fig.** Vertical profiles of chlorophyll-a contents (mg/m^3^) in the water columns of (A) Mejillones Bay, and (B) Antofagasta Bay

**S2 fig.** Vertical profiles of (A) Ammonium (µM); (B) Nitrite (µM); (C) Nitrate; and (D) Phosphate (µM) in the water column of Mejillones Bay.

**S3 fig.** Vertical profiles of (A) Ammonium (µM); (B) Nitrite (µM); (C) Nitrate; and (D) Phosphate (µM) in the water column of Antofagasta Bay.

**S4 fig.** Vertical profiles of chlorin index and TOC (mg/L) in the water columns of Mejillones Bay (A and C), and Antofagasta Bay (B and D).

**S5 fig.** Box-plot diagram showing the δ^13^C mean values of THAA obtained from suspended POM collected in Mejillones and Antofagasta bays. Error bars show the standard error. Black dots represent de values jitter.

**S6 fig.** Spearman rank order correlation in A) Mejillones Bay, and B) Antofagasta Bay. Significant values (p<0.05) are boxed.

**S1 table.** Two samples Mann-Whitney tests (significant values = p<0.05) for temperature, salinity, nitrite (NO_2_ ^-^), nitrate (NO_3_ ^-^), and phosphate (PO_4_ ^3-^) in Mejillones and Antofagasta bays.

